# Roles of signaling compounds and WRKY31 in the defense of *Pinus massoniana* L. against *Dendrolimus punctatus*

**DOI:** 10.1101/2020.10.13.337279

**Authors:** Hu Chen, Ying Hu, Xingxing Liang, Junkang Xie, Huilan Xu, Qunfeng Luo, Zhangqi Yang

## Abstract

*Dendrolimus punctatus* is an important pest affecting Masson pine (*Pinus massoniana* L.) forests and can cause serious economic and ecological losses. WRKY transcription factors play important roles in coping with various environmental stresses. In particular, recent studies have shown that WRKY transcription factors play an important role in plant responses against herbivorous insects. However, the mechanisms underlying the actions of these genes in the defense responses of *P. massoniana* L. are still unclear. Our previous study provided evidence that WRKY may play an important role in the insect resistance of *P. massoniana* L. In this study, application of semiochemicals such as exogenous hormones and Ca^2+^ by spraying increased the concentrations of endogenous hormones, terpenoid synthases, and volatile substances in *P. massoniana* L. and effectively improved its resistance to *D. punctatus*. After analyzing the WRKY family of *P. massoniana* L., the PmWRKY31 gene was selected and studied. Yeast two-hybrid assays showed that the LP8 gene interacted with PmWRKY31. Fluorescence-based quantitative polymerase chain reaction showed that after treatment with exogenous hormones and Ca^2+^, the expression levels of the PmWRKY31 gene, hormonal signal–related genes, and terpene biosynthetic pathway–related genes were significantly increased, whereas the expression of the LP8 gene was decreased. Therefore, the PmWRKY31 and LP8 genes affected downstream gene expression by positively and negatively regulating the hormone signaling pathways, respectively. This result provides theoretical support for the involvement of WRKY transcription factors in the insect resistance of *P. massoniana* L. through their regulation of hormone signaling.

## Introduction

Herbivorous insects are important pests affecting agricultural and forestry production and can result in severe economic and ecological losses. The interaction between plants and insects can activate defense responses in plants[1]. This behavior is the first line of defense for plants. It involves the activation of different signal transduction pathways and downstream chain reactions, and the related transcription factors regulate defense gene transcription to synthesize special defense compounds and initiate defense responses[2].

The jasmonic acid (JA), salicylic acid (SA), ethylene (ET), and Ca^2+^ signaling pathways play important roles in plant defense against insects[3–11]. Silencing of JA biosynthesis–related genes OsHI-LOX, AOS1, and AOS2 significantly reduces the damage of brown planthopper to rice plants and is regulated by transcription factors such as OsERF3, OsWRKY70, and OsWRKY24[7–9,12]. SA plays important roles in inducing insect resistance[13]. JA and SA can mutually induce defensive gene expression, indicating the synergy between the two[14].

Insect feeding induces the expression of defense genes in plants, as well as Ca^2+^ flow and changes in intracellular JA and SA[15]. Ca^2+^ and Ca^2+^-dependent protein kinase regulation is critical for enhancing the resistance of Arabidopsis to *Spodoptera littoralis*[16–17]. In the development of plant resistance, Ca^2+^, SA and JA cross-talk and enhance plant resistance through signaling cascades[18–19].

All plants have their own ‘fragrance’ to repel herbivorous insects, and it is key to determine the genes that regulate certain substances to produce as much of the ‘fragrance’ as possible to ‘reject’ the herbivorous insects from feeding[9]. Plants can generate many secondary metabolites, especially volatile metabolites such as terpenes, to avoid harm from herbivorous insects. In the process of plant defense against herbivorous insects, the production of secondary metabolites is regulated by various hormones, such as JA, SA, and gibberellin (GA)[20]. WRKY transcription factors play important roles in regulating the resistance to insects, diseases, and abiotic stress in plants[21–25]. Among the WRKY transcription factors, WRKY3 and WRKY6 can regulate the insect resistance of tobacco plants and participate in the JA pathway [26]. WRKY genes participate in the mechanism of insect resistance in tomato and *Arabidopsis[27–28]*, Systematic studies of rice WRKY transcription factors in insect defense have shown that they participate in a variety of hormone metabolic pathways enhancing insect resistance in rice[29–30].

*Pinus massoniana* L. is a very important timber species in China, accounting for more than half of the growing stock of forests in South China. It is also the major resin-producing tree species in the world[31]. The insect *D. punctatus* causes severe damage to approximately 130000 hectare of Masson pine forests every year, severely affecting tree growth and forest ecology[32]. Currently, the defense mechanism of *P. massoniana* L. against *D. punctatus* remains unclear. To understand the defensive signaling pathways of *P. massoniana* L., on the basis of a previous transcriptome analysis, we speculated that WRKY transcription factors play an important role in this defense. This study investigated the mechanisms of the WRKY-based defense system by screening their interacting genes, analyzing the hormones they stimulate, and detecting volatile substances in this pine.

## Materials and Methods

### Plant growth and growth conditions

The seeds of *P. massoniana* L. (No. 17-243) were from the F1 generation (Nanning China). Tobacco plants were grown from *Nicotiana benthamiana* seeds. The seeds were stored in a refrigerator at 4 °C until use.

In April 2017, the seeds were sown in yellow soil for germination. When the buds grew to 5 cm tall, they were transplanted into nonwoven bags with a diameter of 12-15 cm. The light medium was formulated with 45-60% peat or coconut chaff, 20-30% carbonized rice chaff, 8-9.5% perlite, 1% calcium superphosphate, and 10-15% peat soil. The buds were planted in the breeding nursery for pine seedlings of the Guangxi Academy of Sciences (Nanning, Guangxi, China). Healthy 1-year-old seedlings of *P. massoniana* L. with good growth, the same height, no insect damage, and no mechanical damage were selected as experimental materials.

### Experimental *D. punctatus*

The *D. punctatus* cocoons for experimental use were collected from the Masson pine orchards (Ningming and Nanning, Guangxi, China) and cultured in an incubator at 26 ± 0.5 °C, under 16 h of light each day, and at relative humidity of 80%. Second-instar larvae were used for experiments.

### Plant treatments

The following treatment solutions were prepared: 75 mg/L abscisic acid (ABA), 75 mg/L ABA + 100 mg/L CaCl_2_, 50 mg/L SA, 50 mg/L SA + 100 mg/L CaCl_2_, 100 mg/L MeJA, and 100 mg/L MeJA +100 mg/L CaCl_2_, 150 mg/L GA, 150 mg/L GA + 100 mg/L CaCl_2_, and 100 mg/L CaCl_2_ (dissolved in 50 mM phosphate buffer, pH 8.0). Distilled water was used as a control. Before use, 0.01 (v/v) Tween-20 was added to the solutions, and the treatment solutions were evenly sprayed on the *P. massoniana* L. seedlings once a day for 5 days at 200 mL/treatment. Ten *P. massoniana* L. seedlings with uniform growth were used for each treatment. Mature needles at the same site were collected 1, 3, and 5 days after the treatment ended. The needles collected were divided into two groups. One group was immediately tested to measure the volatile substances. The other group was immediately stored in liquid nitrogen, transferred to the laboratory, and stored at −80 °C for future use. ABA, SA, JA, and GA (purity> 95%) were purchased from Sigma-Aldrich.

### *D. punctatus* feeding treatment

An insect incubator was used for each *D. punctatus* feeding treatment, and 15 *D. punctatus* of same size were selected for each treatment. On day 3, the *D. punctatus w*ere observed. The effects of different treatments on *D. punctatus* were investigated based on their food intake and growth conditions. The food intake calculation formula for larvae was as follows: daily food intake = (amount of feed input - amount of residual feed) × (1 - water loss rate). The treatments were carried out in net houses.

### RNA extraction and reverse transcription

RNA was extracted according to the instructions of the RNA Isolation Kit for polyphenol- and polysaccharide-rich plants (Tiangen Biotech, Beijing, China). The reverse transcription primer was Oligo(dT)18: 5’-GGCCACGCGTCGACTAG TAC(T)18-3’. The specific cDNA was synthesized according to the instructions of M-MLV reverse transcriptase. After completion, 4 μL of the polymerase chain reaction (PCR) product of each treatment was used for agarose gel electrophoresis, and the cDNA concentration of each treatment was measured using a UV spectrophotometer and then diluted to the same concentration.

### Isolation and characterization of gene expression from cDNA

Based on transcriptome data (transcriptome data uploaded to NCBI GEO, accession number GSE72294.) and protein–protein interactions, the full lengths of the WRKY2, WRKY6, PMWRKY31, and LP8 genes were obtained. Primer 5 software was used to design full-length primers to amplify these genes (Table S1). The PCR products were cloned into the pMD19-T vector (TaKaRa) and sent to Sangon Biotech (Shanghai, China) for sequencing.

### Bioinformatics analysis

WoLFPSORT software was used to predict the subcellular localization of proteins. The amino acid sequences of the proteins were constructed with ClustalX and MEGA4.1 software. The online software of NCBI, SMART, and Motif Scan were used to analyze the functional domains of genes. Protein–protein interactions were predicted using string (https://string-db.org/cgi). Transcriptome data and QuickGO (https://www.ebi.ac.uk/QuickGO/) were used to predict gene function, and the Kyoto Encyclopedia of Genes and Genomes (KEGG) data were used for metabolic pathway analysis.

### Subcellular localization

The constructed pBWA(V)HS-wrky-GLosgfp vector plasmid was transferred into Agrobacterium. After Agrobacterium-coated plates were incubated at 30 °C for 2 days, Agrobacterium was inoculated into 10 mL YEB liquid medium and resuspended in 10 mM MgCl_2_ suspension (containing 120 μM AS), and the optical density measured at a wavelength of 600 nm (OD600) was adjusted to approximately 0.6. The suspension was injected into the epidermis of a tobacco leaf with a 1-mL syringe (needle removed). After injection, the tobacco plants were cultured under low light intensity for 2 days. Next, the tobacco leaves were collected and imaged directly under a laser confocal microscope (FV10-ASW, OLYMPUS, Shenzhen, China). In the subcellular colocalization experiment, except that the nuclear marker and the plasmid vector were simultaneously transferred to Agrobacterium before plating and incubation, all other steps were the same.

### Transgene expression

Sterile tobacco seedlings were induced using mature tobacco embryos. The plasmid pBI121_PmWRKY31 was constructed and transferred to Agrobacterium EHA105 and stored in a −80 °C freezer. The Agrobacterium-mediated transformation of tobacco seedlings followed the steps described by Yu et al.[33]. The cetyltrimethylammonium bromide (CTAB)-based method was used to extract DNA from tobacco seedlings, and primers specific for resistance genes (Table S1) were used to amplify and detect the presence of PmWRKY31 in tobacco seedlings using the conventional PCR method. Transgenic lines were screened from the F3 generation of tobacco plants transduced with PmWRKY31, morphological indicators were observed, and hormones, volatile substances, and resistance were determined.

### Yeast two-hybrid assay

#### Construction of the cDNA library

After RNA extraction, cDNA was synthesized and purified. The purified cDNA was homogenized and further purified. The cDNA was digested using the restriction endonuclease SfiI. After the digested cDNA was passed through CHROMA SPIN-1000-TE columns, an appropriate amount of cDNA was ligated into the pGADT7-SfiI vector (TaKaRa, China) at 12 °C using the DNA ligation kit (O/N linked) and purified to obtain the primary cDNA library, which was electroporated into HST08 competent cells. Ten large Luria-Bertani (LB) agar plates (24.5 × 24.5 cm) were coated with these cells and cultured overnight at 37 °C, and the number of clones obtained after the transformation was monitored.

### Yeast two- and four-hybrid assays

Five micrograms of the bait plasmid was transformed into Y187 yeast, and 100 SD/Leu plates were coated with the yeast and cultured at 30 °C for 3 days. The pGBKT7-PmWRKY31 plasmid vector was constructed, and the bait plasmid was transformed into the Y2Hgold strain to obtain the bait strain. The expression of the exogenous proteins in the bait strain was detected by western blotting.

Two-hybrid screening: Bait-Y2HGold strains were cultured using the streak plate method for 3 days. Colonies were picked and cultured in SD/-Trp broth and mated to the Y187 yeast library. A small amount of suspension was diluted to 1/10, 1/100, 1/1000, and 1/10000, and 100 μL of the diluted suspension was used to coat 100-mm monitoring plates. The suspension was coated onto 50 to 55 SD/-Trp/-Leu/X-a-Gal/Aba plates for two-hybrid screening.

Four-hybrid screening: The blue colonies were counted and inoculated onto SD/-Ade/-His/-Leu/-Trp/X/A plates with a pipette tip and cultured at 30 °C for 5 days. The positive bacterial strain was used as a template for PCR amplification. The AD plasmids in the positive clones were detected, and the amplified products were detected by electrophoresis and sequenced.

### Bimolecular fluorescence complementation (BiFC)

The LP8, WRKY2, WRKY6, and PmWRKY31 genes were separately cloned into the pSPYNE-35S vector. The proteases were isolated from 3-to-4-week-old Arabidopsis plants with robust growth, transfected by the polyethylene glycol (PEG) method, and observed under a confocal microscope (FV10-ASW, OLYMPUS).

### Pull-down assay

Primers for the PmLP8 and PmWRKY31 genes were designed in CmSuite8 software (Table S1). The target gene fragment was amplified by the high-fidelity PrimeSTAR DNA Polymerase with the WRKY plasmid as the template. Five micrograms of the pGEX-4T-1 vector was digested with XhoI and BamHI to recover the target fragment. Ligation was performed according to the manufacturer’s instructions of the ClonExpress II One Step Cloning Kit (Vazyme Biotech). The ligation products were transformed into Stbl3 competent cells and screened on LB plates containing kanamycin and ampicillin antibiotics (100 μg/mL). Positive clones were confirmed by sequencing.

The fusion protein was subjected to prokaryotic expression, pull-down, western blot detection, electrophoresis, transfer onto membranes, antibody incubation, and exposure according to the manufacturers’ instructions. GST antibody and HIS antibody were purchased from TRANS (Shenzhen, China), and HRP-labeled goat anti-mouse IgG was purchased from CWBiotech (Beijing, China).

### Real-time fluorescence-based quantitative PCR

Key genes involved in metabolic pathways, such as ABA, GA, JA, SA, and terpene biosynthetic pathways, which might interact with WRKY genes, were selected. Primer 5 software was used to design primers for fluorescence-based quantitative PCR, and the cyclophilin (CYP) gene was used as a reference gene (Table S1) [34]. The LightCycler 480II PCR system was programmed according to the instructions of the SYBR Premix Ex *Taq* II (Perfect Real Time) kit (TaKaRa, China) to conduct fluorescence-based quantitative PCR. All experiments were run three times. The relative expression level was calculated according to the 2^-ΔΔCt^ method[35], and Microsoft Excel was used for plotting.

### Determination of volatiles substances

Volatile substances were determined by a SCION single-quadrupole (SQ) and triple-quadrupole (TQ) gas chromatography (GC)–mass spectrometry (MS) system. Each treatment (0.5 g) was placed in a 10-mL-headspace bottle, and an appropriate amount of anhydrous sodium sulfate was added. After an aged solid-phase microextraction fiber was inserted into the bottle, the bottle was sealed and put into a 75 °C thermostat bath for 15 min. The extract was subjected to GC-MS analysis. Each experiment was run three times.

### Determination of semiochemicals

First, 0.1 g of sample was ground in liquid nitrogen, added to 1 mL methanol solution (methanol:water:formic acid = 75:20:5). After 16 hours of extraction in darkness, the supernatant was collected by centrifugation. The above steps were repeated once, and the supernatant was collected and combined with the previously obtained supernatant. The combined supernatant was concentrated and evaporated at 35 °C until there was no residual methanol (changed color). Then, 500 microliters of ethyl acetate was added to the remaining aqueous phase for extraction, and the upper, ester phase was taken. This step was repeated two times, and the obtained ester phases were combined. The combined ester phase was concentrated and evaporated at 35 °C till dry. The precipitate was dissolved in 200 μL of methanol, filtered through a 0.22-nm organic membrane, and tested by a liquid chromatography (LC)–MS system (6460 Triple Quad LC/MS, Agilent, USA) with a C18 column (2.1 mm × 100 mm, 1.9 μm). According to the plotted standard curve and the peak area of the substance in the sample tested, the concentration of the substance in the sample was calculated.

### Detection of terpene synthases (TPSs)

A total of 0.1 g mixed sample was added into 900 μL of phosphate-buffered saline (1:9 weight:volume ratio) and fully ground to homogenate on ice. After the homogenate was centrifuged at 5000 × g for 5-10 min, the supernatant was taken for detection. A plant TPS ELISA kit for (Shanghai, China) was used for the detection of TPSs in different samples according to the manufacturer’s instructions. The microplate reader was purchased from Epoch (BioTek, USA).

## Results and analysis

### The role of WRKY transcription factors in insect resistance

Three WRKY genes were highly expressed in insect-resistant Masson’s pine varieties (Fig. 1a). According to the cDNA sequences of the three genes obtained from the transcriptome data, they were PmWRKY2, PmWRKY6, and PmWRKY31, encoding 667, 575, and 642 amino acids, respectively (Table S2). All three genes had the typical WRKY domains of the WRKY family (Fig.1b, Fig. 1c, Table S2). WRKY2 was successfully annotated in the KEGG Orthology (KO), indicating that WRKY2 is involved in plant pathogen defense (Fig. 1d). Therefore, we hypothesized that WRKY2 is involved in the defense of *P. massoniana* L. against herbivorous insects.

**Fig 1.**
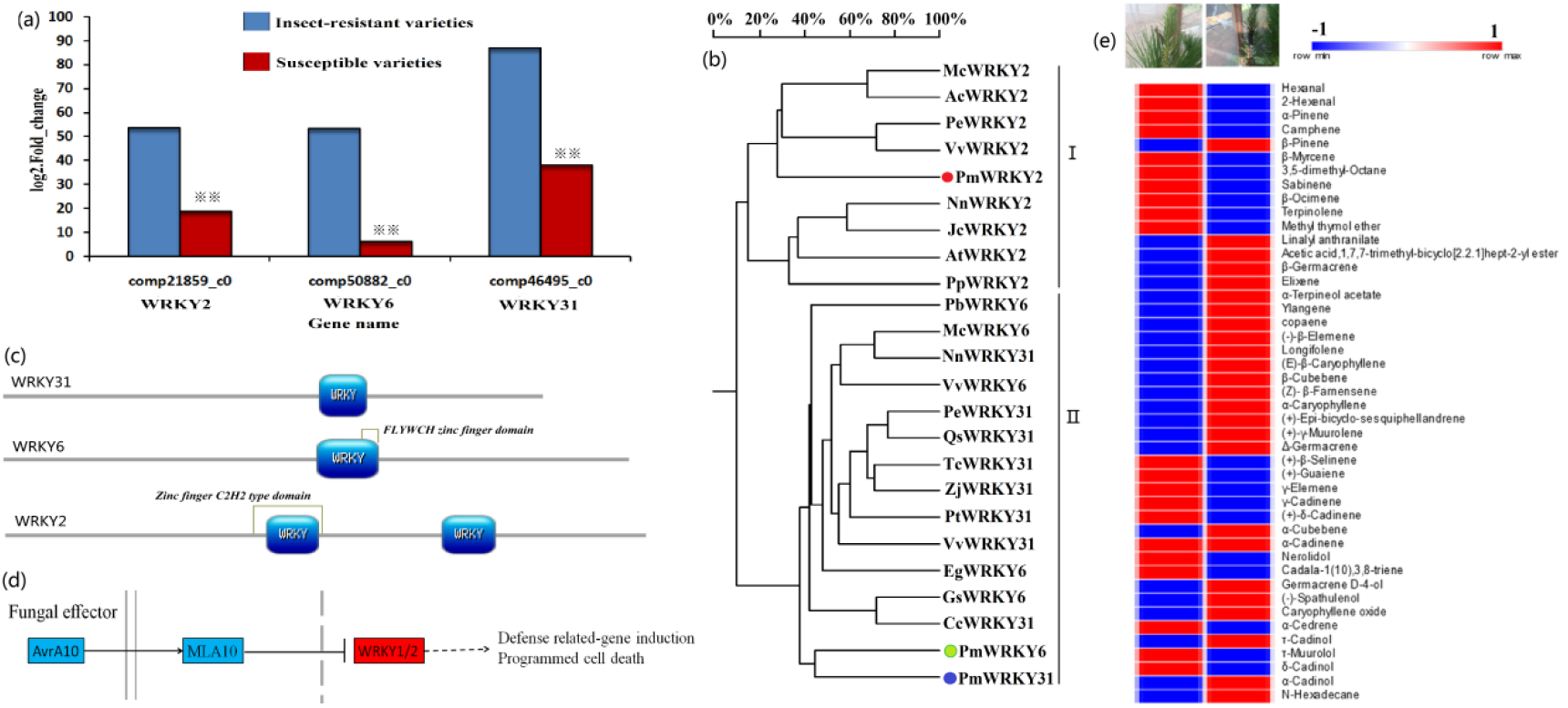
WRKY gene in insect resistant and bioinformation analysis. a: three WRKY transcriptomes obtained from 511 differeniationly genes in High throughput sequencing analysis of transcriptome of insect resistant varieties Vs susceptible varieties. The data is log2.Fold_change of insect resistant varieties Vs susceptible varieties, and each sample transcriptome replicated three times. b: The analysis of WRKY polygenetic tree. Pm, *Pinus massoniana* L.; Nn, Nelumbo nucifera; Vv, Vitis vinifera; Qs, Quercus suber; Pt, *Populus trichocarpa*; Zj, *Ziziphus jujube*; Cc, *Cajanus cajan*; Jr, *Juglans regia*; Tc, *Theobroma cacao*; Mc, *Macleaya cordata*; Ac, *Aquilegia coerulea*; Pp, *Physcomitrella patens*; At, *Amborella trichopoda*; Jc, *Jatropha curcas*; Pe, *Populus euphratica*; Gs, *Glycine soja*; Pb, *Pyrus* x *bretschneideri*; Eg, *Elaeis guineensis.* The number of different WRKYs are NnWRKY31 (XP_010252466.1); VvWRKY31 (XP_002269696.2); QsWRKY31 (XP_023921697.1); PtWRKY31 (XP_002321134.3); ZjWRKY31 (XP_015877768.1); CcWRKY31 (XP_020234210.1); JrWRKY31 (XP_018811738.1); TcWRKY31 (EOX93243.1); NnWRKY2 (XP_010270167.1); PpWRKY2 (XP_024368161.1); AtWRKY2 (XP_006836767.1);VvWRKY2 (CBI39865.3); JcWRKY2 (XP_012070967.1); ZjWRKY6 (XP_015877768.1); QsWRKY6 (XP_023921697.1); GsWRKY6 (KHN36523.1); NnWRKY6 (XP_010252466.1); PbWRKY6 (XP_018502314.1); McWRKY6 (OVA03405.1); EgWRKY6 (XP_010926185.1); PeWRKY6 (XP_011047241.1); VvWRKY6 (XP_002263115.1). c: Prediction of NCBI blasts, SMART and Motif Scan online software for functional domain analysis of four genes WRLY2, WRKY6, WRKY31, and Lp8 with Prosite software (https://prosite.expasy.org/mydomains) were mapped. Colored sections are the main functional domains of the genes. d: The annotation and signaling pathways of WRKY2 gene using KEGG database. b Results of needles volatile substances of insect resistant varieties and susceptible varieties in May. The results were analyzed by GC and GC-MS (HP6890 gas chromatograph ((Hewlett-Packard Company, USA), GCMS-QP5050A gas chromatography-mass spectrometry (Shimadzu Corporation, Japan)). The sample was repeated three times.

Seventy-four volatile substances were detected, and 45 of them were successfully identified. Among the identified volatile substances, 30 were terpenes. α-Pinene and β-pinene showed opposite abundance patterns (Fig. 1e). TPSs and hormones play roles in insect resistance (Sakamoto et al., 2004; Eric et al., 2010; Hu et al., 2015; Liu et al., 2016). Whether both WRKYs and TPSs participate in the insect resistance of *P. massoniana* L. and their relationships with semiochemicals needed to be clarified.

### Screening of PmWRKY31 transcription factor interaction genes

To further investigate the function of PmWRKY31 in the insect resistance of *P. massoniana* L., we constructed the cDNA library (Fig. S1) and the two-hybrid bait system (Fig. S2). The bait strains did not self-activate and had no toxicity (Fig. S3a). They were highly expressed in the positive control according to the western blot (Fig. 2a). Two colonies (Fig. S3b-d) were obtained by two-hybrid screening and four-hybrid screening (Fig. 2b). The positive control and negative control showed the expected results (Fig. 2c). Sequencing of the PCR amplification product revealed the calcium-binding protein LP8 (Fig. 2d). This gene contained three EF-hand calcium-binding domains and a secreted protein acidic and rich-in-cysteine Ca^2+^-binding region (Fig. 2e).

**Fig 2.**
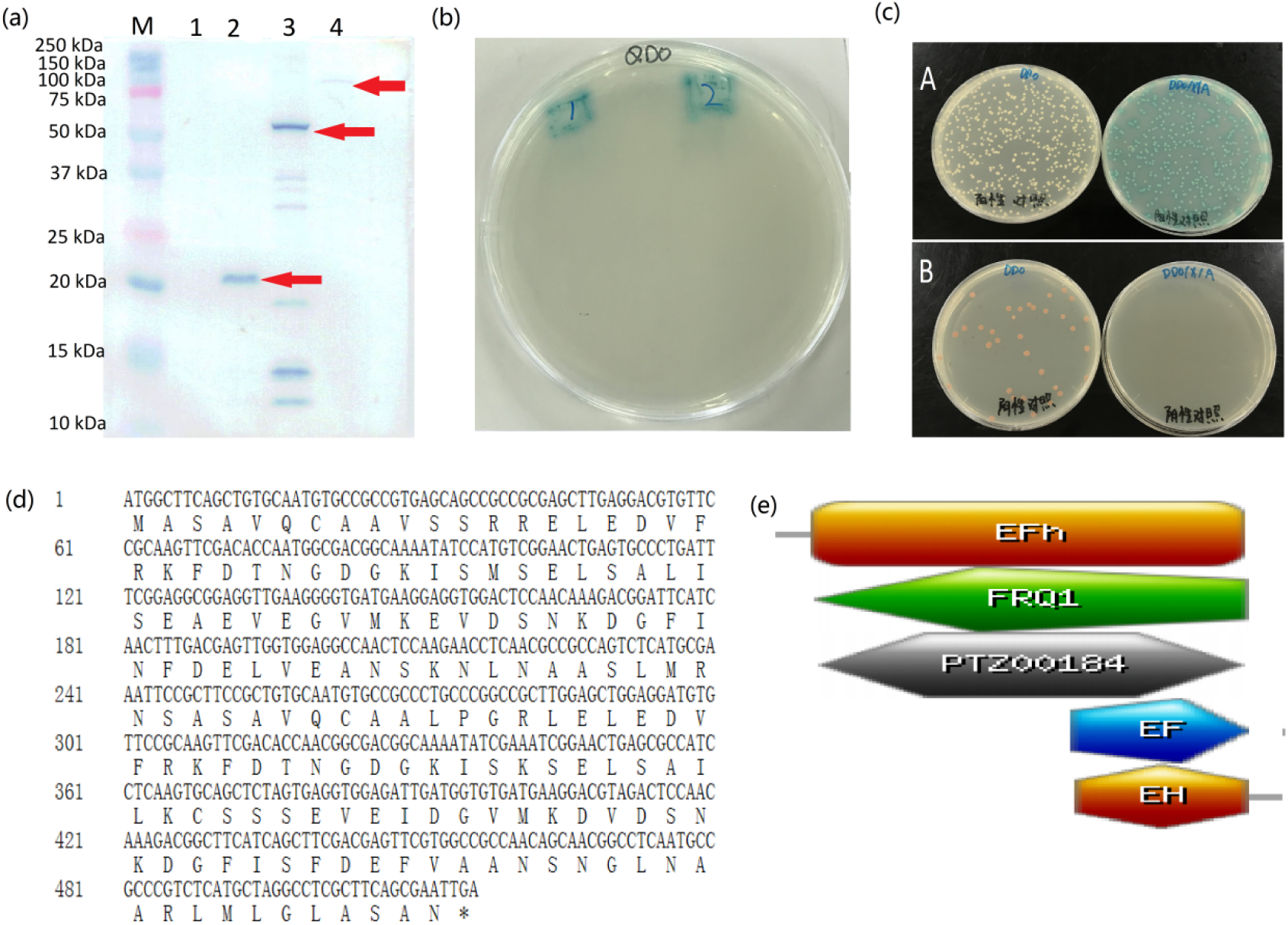
Two-hybrid screening of PmWRKY31 interactional gene and analysis of interactional gene. a: Detection of exogenous protein expression of two-hybrid bait. Validation results of Western blot of Bait expression using Gal4-BD monoclonal antibody in yeast two-hybrid experiment. Lane 1: Y2HGold strain without plasmid (negative control); lane 2: Y2HGold strain with pGBKT7-BD plasmid (GAL4-BD protein size: 22 KDa); lane 3: Y2HGold strain with pGBKT7-53 plasmid (protein size: 53KDa); lane 4: Y2HGold strain with PmWRKY31Bait plasmid (approximate protein size: 92KDa). b: Positive strains obtained from screening. c: positive and negative control for the two-hybrid screening, A: positive control. Y2Hgold strains with pGBKT7-53 plasmid + pGADT7-T plasmid were coated to DDO plates and DDO/X/A plates; B: Negative control. Y2Hgold strains with pGBKT7-Lam plasmid + pGADT7-T plasmid were coated to DDO plate and DDO/X/A plate. D: Nucleic acid sequence and amino acid sequence information of the Lp8 gene. E: Analysis of the main functional domains of the Lp8 gene.

### Confirmation of the interaction between PmWRKYY31 and PmLP8

To further confirm the interaction between PmWRKY31 and LP8, we performed pull-down experiments (Fig. 3a). We also used BiFC technology to further confirm the protein–protein interactions between LP8 and the three WRKY genes of *P. massoniana* L. (Fig. S4). The results showed that LP8 interacted with WRKY2, WRKY6, and WRKY 31 in the nuclei of plant cells (Fig. 3b). Subcellular localization of WRKY31 also indicated that WRKY31 was located in the nucleus (Fig. 3c, d).

**Fig 3.**
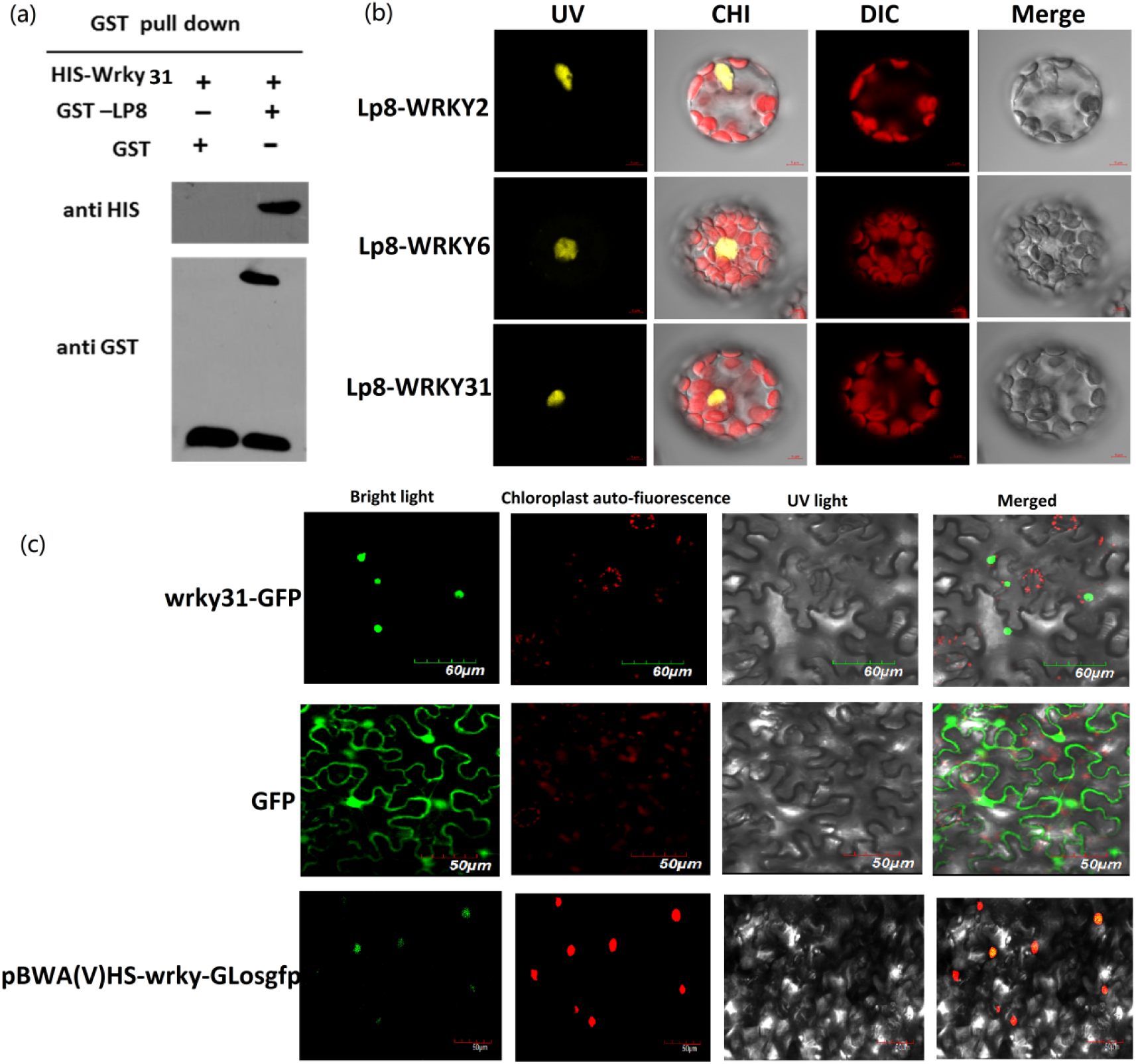
Interaction between PmWRKY31 and Lp8 *in vitro* and *in vivo*. a: Pull down experiments of PmWRKY31 and Lp8 *in vitro*. Constructed a GST prokaryotic expression vector for the bait Lp8 protein as well as his prokaryotic expression vector of the prey PmWRKY31 protein, respectively. Western blot was performed after adding loading buffer to GST-LP8 and His-Wrky fusion proteins to verify the normal expression of the fusion proteins. The GST protein, GST-LP8 protein with GST resin were incubated with His-Wrky protein overnight and eluted with reduced glutathione the next day. The following day, the elution was performed with reduced glutathione. Western blot was performed after appropriate amount of the eluate was treated with loading buffer. b: BiFC validated interactions of PmLP8 and PmWRKY2, PmWRKY6. PmWRKY31. Construction of pSPYNE-35S-lp8, pSPYCE-35S-wrky2, pSPYCE-35S-wrky6, pSPYCE-35S-wrky31 vectors were used for BiFC with *Arabidopsis thaliana*. From left to right, the pictures were yellow fluorescence channel, red fluorescence channel, bright field, and superimposed map. c: localization of PmWRKY31 gene subcellular. Constructed vector plasmids were transfered into Agrobacterium, and tobacco plants in good growth condition were selected. 1 mL syringe without gun tip was used to inject from the lower epidermis of tobacco leaves and then labeled; plants were incubated in weak light for 2 days after injection. Tobacco leaves were taken, observed and photographed with confocal laser microscope (Olympus FV1000, excitation light: 480, emitting light: 510). From left to right, the pictures were bright light, Chloroplast auto-fiuorescence.

### PmWRKY31 improves insect resistance by negatively regulating LP8 and participating in JA signaling

To confirm whether PmWRKY31 regulates these signaling pathways, we examined the semiochemicals, gene expression, terpenes, and feeding characteristics of *D. punctatus* under treatment with these substances.

JA under Ca^2+^ treatment did not differ from the control level on day 1, but JA under the other treatments was significantly different from the control at all times (P<0.01). The JA and JA + Ca treatments significantly improved the JA concentration (Fig. 4a). Moreover, the JA + Ca treatment significantly increased the TPS concentration and increased the concentrations of a variety of volatile substances (Fig. 4b). Different treatments increased many volatile substances in the needles, such as caryophyllene and β-pinene (Fig. 4c). The *D. punctatus* feeding experiment showed that after treatments, the food intake and excrement of *D. punctatus* significantly decreased. Particularly, on day 3 of JA treatment, 64% of *D. punctatus* had died, and the weight of the surviving *D. punctatus* had decreased sharply (Fig. 4d).

**Fig 4.**
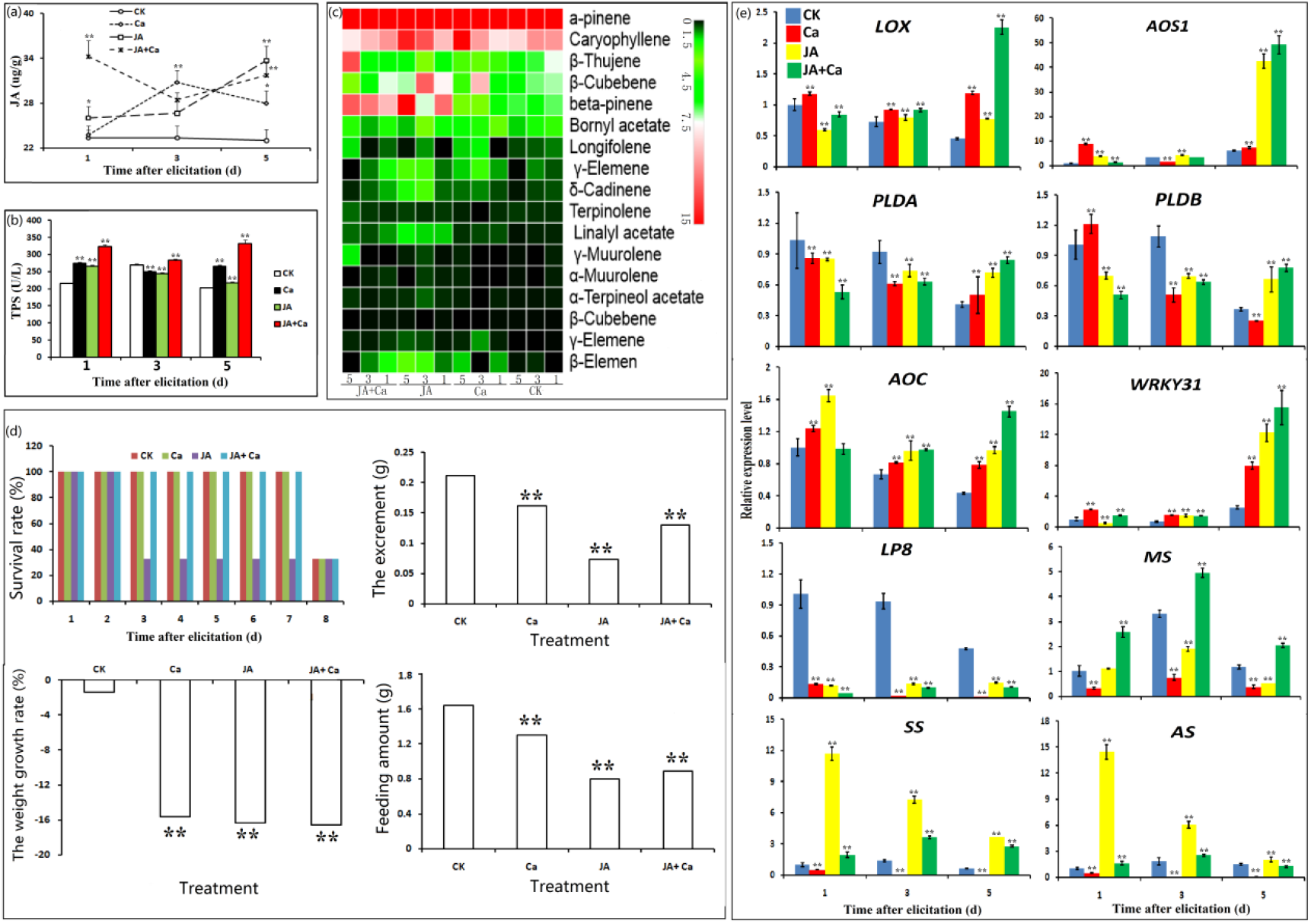
Defense reaction of JA pathway with PmWRKY31. a: JA content in needles in different treatments. b: content of terpene synthase in needles in different treatments. c: Continuous variation of volatile matter content of needles in different treatments. d: Effects of different treatments on feeding and excretion of *Dendrolimus*. e: Expression patterns of terpene synthase genes, key genes of the JA pathway, PmWRKY31 and Lp8 in different treatments. Each sample was repeated 3 times, * p< 0.05, ** p< 0.01, Student’ s t-tests.

To further investigate whether WRKY transcription factors are involved in the insect resistance of *P. massoniana* L. and its responses to different hormone signaling pathways, we performed real-time fluorescence-based quantitative PCR.

PmWRKY31 expression was significantly induced under different treatments, especially on day 5, while the expression of the LP8 gene was downregulated under different treatments. We also examined the expression of five JA biosynthesis–related genes, LOX, AOS1, AOC, PLDA, and PLDB, under different treatments. As shown in Fig. 4, LOX, AOS1, and AOC were significantly increased at the early stage of treatment, while PLDA and PLDB only increased at the late stage of treatment. Treatment with Ca alone did not improve the expression of monoterpene, sesquiterpene, or diterpene genes in the volatile substances. JA treatment increased the expression of SS and AS, while JA + Ca treatment had the most significant improvement on the expression level of MS (Fig. 4e). We speculated that the PmPMWRKY31 gene improves the insect resistance of *P. massoniana* L. by negatively regulating the Ca^2+^ signaling pathway and LP8 activity and participating in the JA metabolic pathway.

### PmWRKY31 improves insect resistance by participating in GA signaling

We further investigated the effect of exogenous GA and Ca treatments on GA biosynthesis. The results showed that on the 5th day after treatment, the TPS concentrations of the three treatments were significantly higher than those of the control (Fig. 5a), indicating that exogenous GA and Ca participated in the biosynthesis of terpenes and significantly improved the concentrations of caryophyllene, β-cubebene, and γ-elemene (Fig. 5c). After treatment, the food intake and weight of *D. punctatus* were significantly lower than those in the control group (Fig. 5d). This indicated that the treatments prevented *D. punctatus* from feeding on *P. massoniana* L. Ca treatment did not significantly increase the GA concentration, while the GA and GA + Ca treatments did (Fig. 5b). At the same time, the expression level of LP8 was significantly decreased after different treatments, and the expression level of PmWRKY31 under Ca treatment was the highest (Fig. 5e). This result indicated that PmWRKY31 positively regulated the GA pathway and LP8 negatively regulated the GA pathway. In addition, exogenous treatments, especially GA treatment, positively regulated terpene biosynthetic pathways. The genes involved in the GA biosynthetic pathway, except for GGPS, were all downregulated (Fig. 5e).

**Fig 5.**
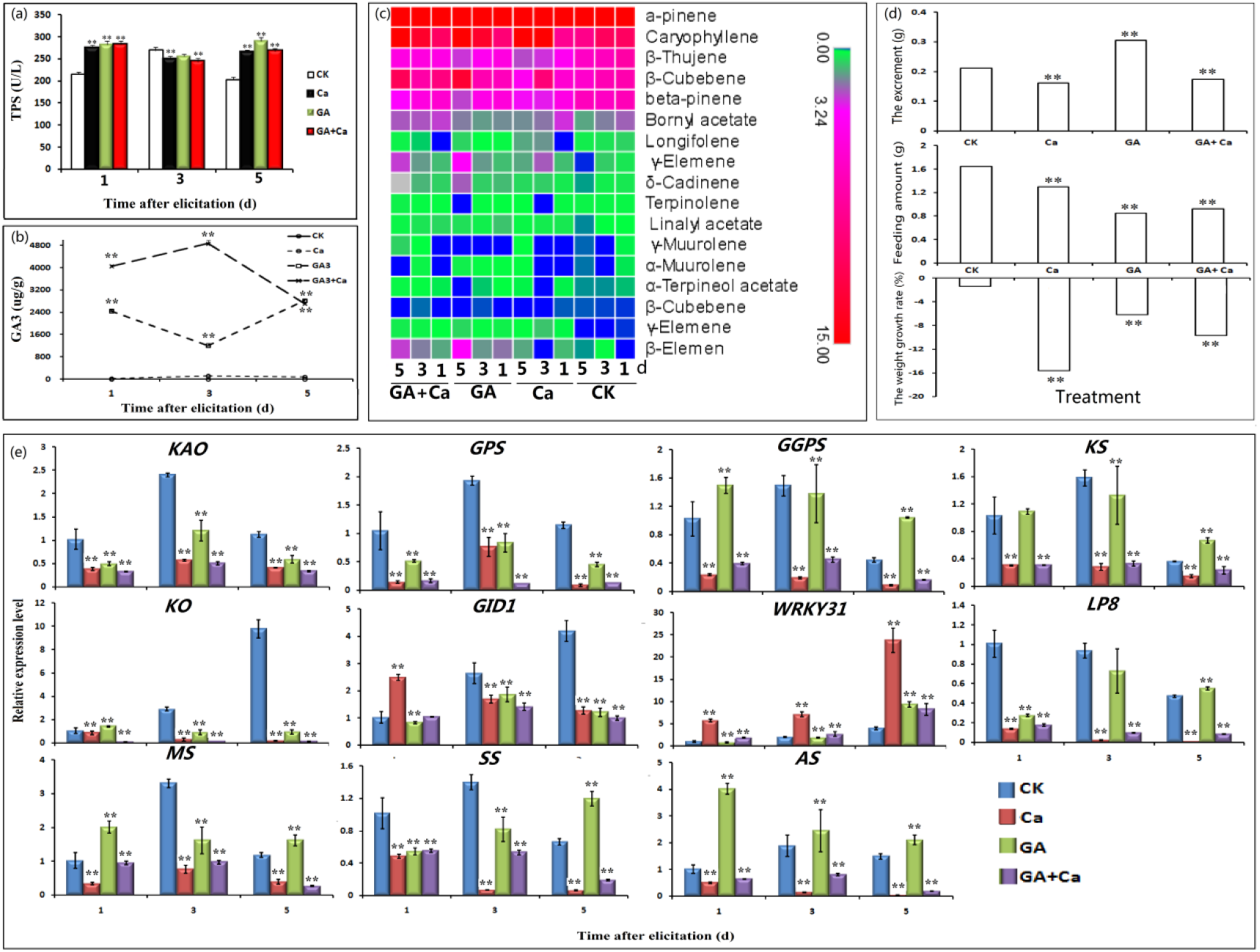
Defense reaction of of GA pathway with PmWRKY31. a: GA content in needles in different treatments. b: content of terpene synthase in needles in different treatments. c: Continuous variation of volatile matter content of needles in different treatments. d: Effects of different treatments on feeding and excretion of *Dendrolimus*. e: Expression patterns of terpene synthase genes, key genes of the GA pathway, PmWRKY31 and Lp8 in different treatments. Each sample was repeated 3 times, * p< 0.05, ** p< 0.01, Student’s t-test.

**Fig 6.**
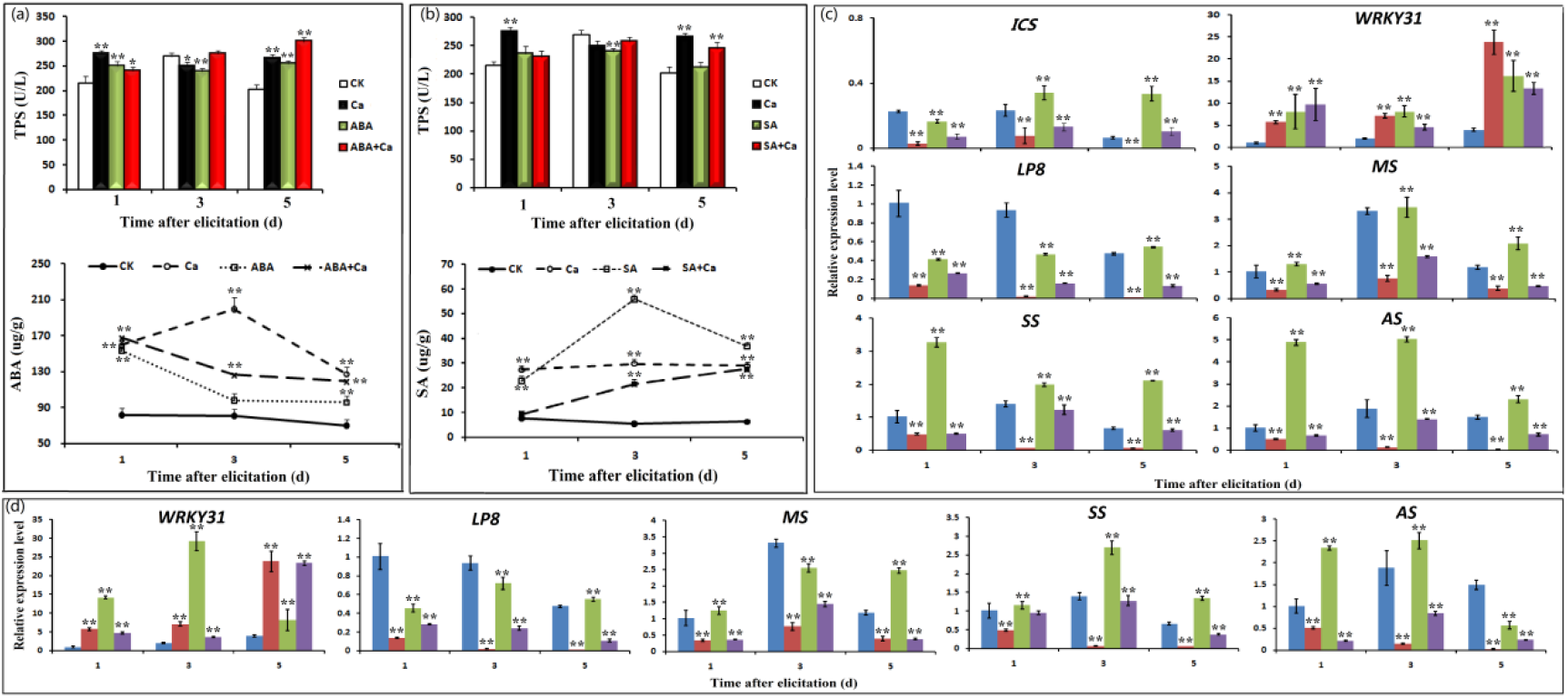
Defense reaction of ABA and SA pathways with PmWRKY31. a: ABA and GA contents in needles in different treatments. b: content of terpene synthase in needles in different treatments. c: Continuous variation of volatile matter content of needles in different treatments. d: Effects of different treatments on feeding and excretion of *Dendrolimus*. e: Expression patterns of terpene synthase genes, key genes of the ABA and SA pathways, PmWRKY31 and Lp8 in different treatments. Each sample was repeated 3 times, * p< 0.05, ** p< 0.01, Student’s t-test.

### PmWRKY31 improves insect resistance by participating in SA and ABA signaling pathways

Under different treatments, the expression level of PmWRKY31 was significantly higher than it was in the control (Fig. 5c, Fig. 5d). The expression of PmLP8 was downregulated under both JA and SA treatments (Fig. 5c, Fig. 5d). Overall, SA treatment alone and ABA treatment alone significantly increased the expression levels of TPS biosynthesis-related genes in the SA and ABA pathways (Fig. 5c, Fig. 5d). However, Ca + ABA treatment significantly increased the TPS level, whereas the application of Ca alone significantly increased the TPS level in the SA pathway. In addition, different treatments had different influences on the concentrations of ABA and SA: Ca treatment significantly increased ABA, and SA treatment worked best at increasing SA (Fig. 5a, Fig. 5b). The substances affected by the four hormone pathways were basically the same. The only difference was that exogenous SA and Ca treatments significantly increased the concentration of bicyclo[4,4,0]dec-1-ene, 2-isopropyl-5-methyl-9-methylene (Fig. S5).

### Verification of the function of the PmWRKY31 gene

To further study the function of the PmWRKY31 gene, we conducted overexpression experiments. The height and root length of PmWRKY31-overexpressing plants were significantly higher than those of the control, and the number of lateral roots, lateral root length, leaf number, and leaf size were significantly lower than those of the control (Fig. 7a). Although the PmWRKY31-overexpressing plants were smaller than the wild-type plants, their resistance to tobacco anthracnose (Fig. 7b) and drought (Fig. 7c) was significantly stronger than that of the wild-type plants. Similarly, the resistance of PmWRKY31-overexpressing tobacco plants against *Helicoverpa assulta* Guenee was also significantly improved. We tried to feed the tobacco leaves of the PmWRKY31-overexpressing and wild-type tobacco plants to *D. punctatus*, but the insect accepted neither of them. In summary, the above experiments further confirm that the PmWRKY31 gene plays a key role in insect resistance as well as resistance to diseases and abiotic stresses.

**Fig 7.**
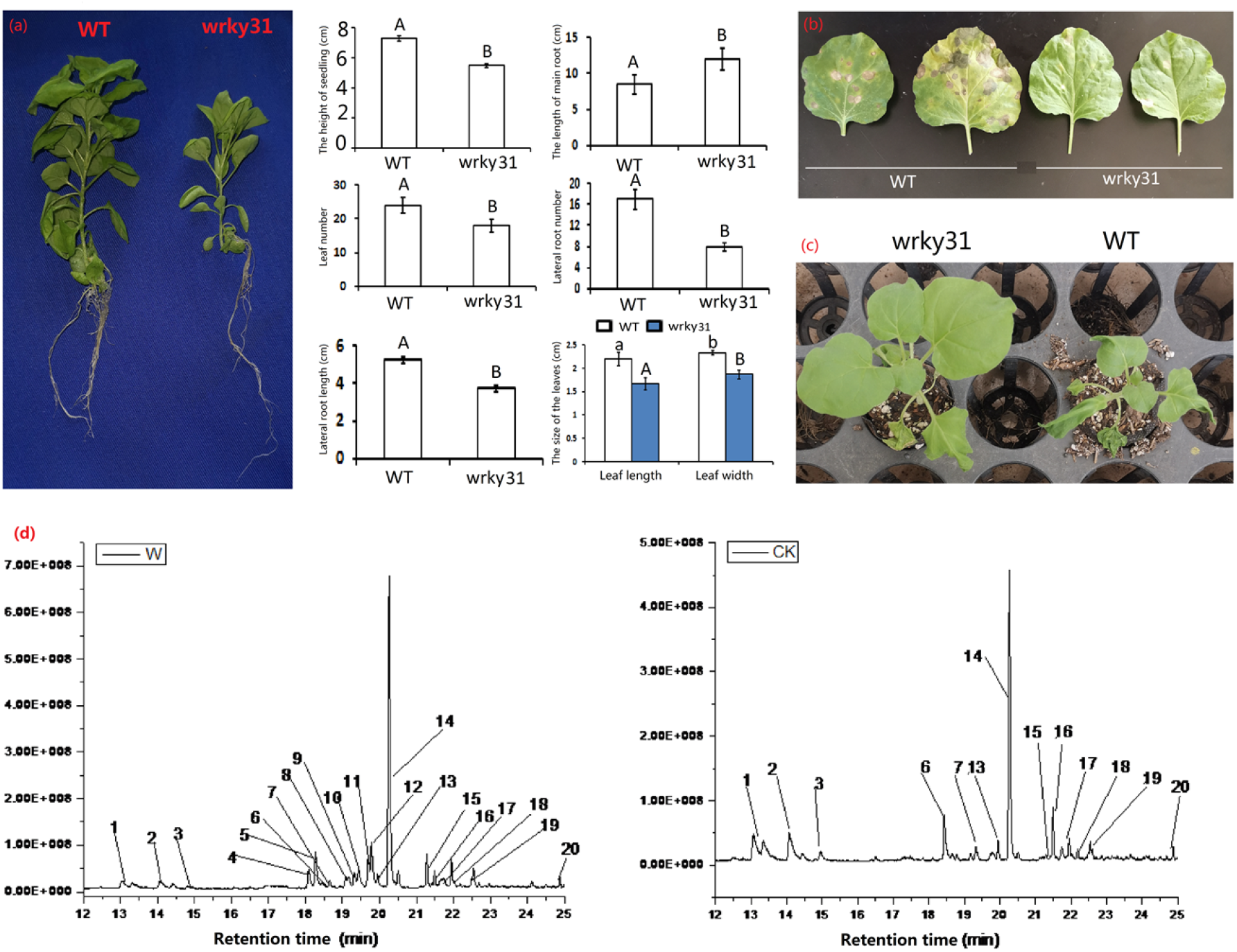
Verification of PmWRKY31 transgene function. a: Effects of transgene on tobacco plants. b: Effects of transgene on disease resistance. c: Effects of transgene on drought tolerance. d: Effect of transgene on volatiles. e: Effect of transgene on hormone content of the plant.

Further analysis of the volatile substances showed that after PmWRKY31 overexpression, the concentrations of more than 10 volatile substances, especially longifolene, changed (Fig. 7d). This result indicated that the changes in terpene concentrations were consistent with the changes in expression levels of TPS genes. We also analyzed the changes in hormone levels. The results showed that the JA and ABA concentrations of PmWRKY31-overexpressing plants were both higher than those of wild-type plants. This indicates that the PmWRKY31 gene regulates the expression of hormone signaling genes by interacting with the PmLP8 gene, thereby increasing the terpene concentrations to improve insect resistance. We also attempted to determine the Ca^2+^ concentrations in needles of *P. massoniana* L. and tobacco leaves, but the experimental data were not ideal.

## Discussion

### Effect of PmWRKY31 overexpression on plant type

Many phenotypes of the PmWRKY31-overexpressing plants were less desirable than those of wild-type plants. In previous studies, OsWRKY28-overexpressing rice plants became dwarf but had no changes in other phenotypes[29,36]. In our study, the situation was different-morphological changes were observed. The causative mechanism is unclear and needs further investigation.

### PmWRKYs are key regulators of insect feeding–induced defense responses

After screening, we obtained three WRKY genes that were associated with the insect resistance of *P. massoniana* L. All of them contained the specific and conserved WRKY domains of the WRKY family and proteins that specifically bind to W-boxes [37]. Clustering analysis can well classify the WRKY genes according to the characteristics of their zinc finger domains[38].

The results of this study all indicated that PmWRKY proteins play important roles in the insect resistance of *P. massoniana* L. First, PmWRKY were induced by JA, GA, ABA, SA and Ca treatments. Second, overexpression of PmWRKY changed the concentrations of JA, GA, ABA, SA, and volatile substances in tobacco plants and improved their insect resistance, disease resistance, and drought resistance. Third, PmWRKY interacted with the Ca^2+^ signaling protein LP8 and changed hormone and TPS concentrations.

### Possible mechanism of the interaction between PmWRKYY31 and Ca^2+^ signaling

Ca^2+^ signaling is involved in plant defense against herbivorous insects, and insect feeding can activate Ca^2+^ signaling[39–41]. However, in the interaction between *D. punctatus* and *P. massoniana* L., the relationship between the Ca^2+^ signaling and the defense response of *P. massoniana* L. induced by *D. punctatus* is still unknown. We found that the addition of exogenous Ca increased the concentrations of TPSs, hormones, and volatile substances in *P. massoniana* L. plants and significantly upregulated PmWRKY31. However, LP8 was downregulated. Therefore, we hypothesized that PmWRKY31 downregulated LP8 to improve the resistance of *P. massoniana* L. against *D. punctatus*. The downregulation of the LP8 gene has been found in a study on the defense of *P. massoniana* L. against *Bursaphelenchus xylophilus[42].* Ca^2+^ phosphorylation regulates the JAV1–JAZ8–WRKY51 network to improve the insect resistance of plants[41]. The Ca^2+^ in chloroplasts can positively regulate MPK3/MPK6 activity[43]. The calcium-dependent protein kinases and mitogen-activated protein kinases (MAPKs) can positively regulate the pathogen defense of *Arabidopsis[44].* WRKY proteins can regulate MAPK genes to improve the insect resistance of rice[7,29,45–46]. In this study, we demonstrated that PmWRKY31 and LP8 can interact both in vivo and in vitro through yeast two-hybrid, BiFC, and pull-down assays and that this interaction played an important role in the responses of *P. massoniana* L. to a herbivorous insect.

### PmWRKY31 plays an important regulatory role in the insect resistance mechanism of *P. massoniana* L. against herbivorous insects

WRKY transcription factors play key roles in hormone signaling pathways and in the regulation of plant resistance genes[26,38,47–49]. We found that application of exogenous JA, SA, ABA, or GA by spraying increased the JA, SA, ABA, or GA concentrations in needles of *P. massoniana* L. and significantly upregulated the PmWRKY31 gene. This result indicates that the PmWRKY31 gene improved the insect resistance of *P. massoniana* L by participating in the JA, SA, ABA, and GA signaling pathways. In addition, the JA and ABA concentrations significantly increased in PmWRKY31-overexpressing tobacco plants. Overexpression of OsWRKY13 and OsWRKY30 in rice enhances the resistance of rice plants to leaf blight and rice blast [50–51]. OsWRKY53 can positively regulate SA biosynthesis[29]. OsWRKY13 can activate ICS1, a key enzyme in SA biosynthesis, and overexpression of OsWRKY13 can induce high accumulation of SA[50]. ThWRKY4 can increase the tolerance of ABA-treated *Tamarix hispida[52].* Based on the above evidence and our experimental results, it can be concluded that the targets of PmWRKY31 could be JA, GA, and SA biosynthesis-related genes, including LOX/AOS1/AOC[1,12], and ICS[53–54]. However, whether PmWRKY31 directly or indirectly binds to the W-box of these gene promoters remains unclear.

### PmWRKY31 plays an important regulatory role in the biosynthesis of terpenes for the insect resistance of *P. massoniana* L

TPSs play a role in insect resistance[55–56]. They are also involved in the biosynthesis of phytoalexin and the regulation of some hormonal substances in the plant defense responses[57]. TPS genes have been cloned from more than 40 species of plants [58].

In the constitutive and induced plant defenses against herbivorous insects, the biosynthesis of terpenes is regulated by a variety of hormones (including endogenous and exogenous hormones). The defensive metabolites of GA and diterpene in rice plants have been studied in detail, and the results indicate that a GAOsCPS1 gene downstream of the GA pathway may be involved in the biosynthesis of terpenes [55]. SA can significantly improve the insect resistance of plants[59–60]. The WRKY1 gene can bind to the W-box of the CAD1-A promoter, activate the expression of CAD1-A, and play an important regulatory role in secondary metabolism. It can also interact with JA and GA signaling molecules to coordinate the biosynthesis and volatilization of terpenes[20]. Our study showed that treatment with one or more of JA, GA, SA, ABA, and Ca^2+^ increased the concentrations of TPSs and volatile terpenes and the expression of monoterpene synthase (MS), sesquiterpene synthase (SS), and diterpene synthase (AS) genes. Overexpression of PmWRKY31 significantly increased the concentration of sesquiterpenes. Therefore, PmWRKY31 might indirectly affect the concentration of terpenes in plants, thereby improving the insect resistance of plants.

### Conclusions

In summary, treatments with exogenous semiochemicals increase the concentrations of endogenous hormones and TPSs of *P. massoniana* L., strongly induce the expression of PmWRKY31, but inhibit the expression of LP8. PmWRYK31 interacts with LP8 and regulates the expression of JA, GA, and SA genes, thereby promoting the expression of TPS genes and increasing the concentrations of volatile substances such as terpenes (Fig. 8). All of the above improve the resistance of *P. massoniana* L. to *D. punctatus.*

**Fig 8.**
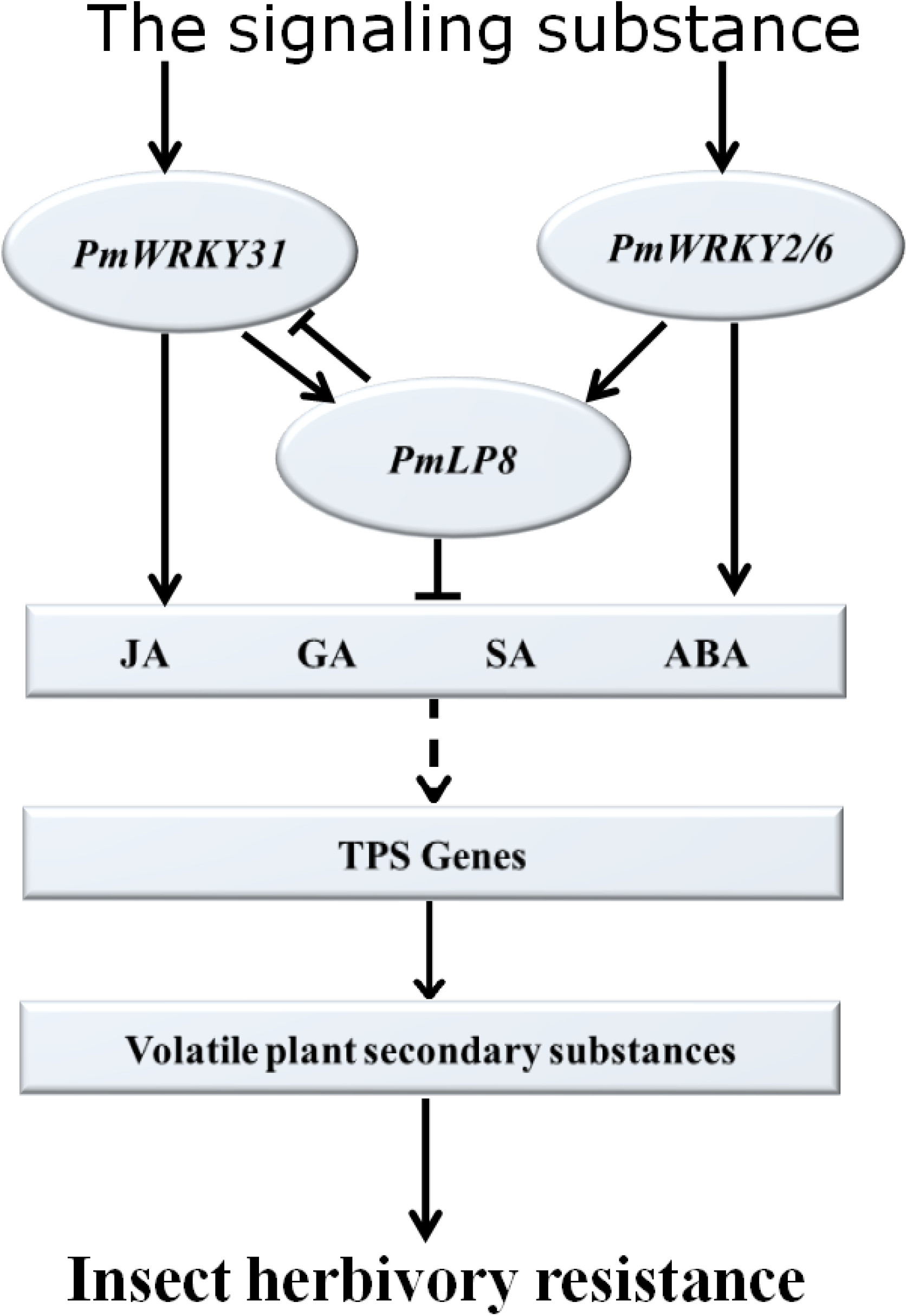
Preliminary model of improvement of insect resistance in *Pinus massoniana* by regulation of hormone signaling pathway of PmWRKY31 and LP8 genes interaction. The application of exogenous signal substances was able to rapidly initiate the expression of PmWRKY31, PmWRKY2 and PmWRKY6 genes, and increase the downstream hormone signals and key gene responses of the terpene synthesis pathway by regulating the LP8 gene, thereby increasing the content of endogenous JA, GA, SA, ABA as well as terpene synthase and volatiles in *Pinus massoniana* to promote the ability to resist pine caterpillars.

## Supplementary Data

Supplementary Data for this article are available at PLOS ONE online.

**Fig S1. Production of cDNA library**

a: RNA extraction. 300 ng of RNA was taken for 1.5% agarose gel electrophoresis to check the quality of RNA. b: Construction of cDNA library. cDNA was synthesized by using Clontech’s SMART cDNA Library Construction Kit and Advantage 2 PCR Kit. 3 μL cDNA was subjected to 1.5% agarose gel electrophoresis. c: Homogenization of cDNA synthesis. PCR was performed by using Normalization Kit and the Advantage 2 PCR Kit amplify to obtain homogenized cDNA, and 1 uL cDNA was taken for 1.5% agarose gel electrophoresis. d: Removal of short fragment. The short fragment of enzyme was removed using CHROMA SPIN-1000-TE. 1 μL cDNA was subjected to 1.5% agarose gel electrophoresis after cleaned up by PCI/CI, refined in ethanol, and solubilized in dH2O. e: Measurement of primary library volume. Amp-resistant LB plates were coated with transfer solution and incubated overnight at 37°C; primary libraries were counted by the number of colonies grown on the plates. 1 library: > approximately 4.4×10^6^cfu; 2 libraries: > approximately 2.8×10^6^cfu; 3 libraries: > approximately 4.6×10^6^cfu. 1: Total RNA of *Pinus massoniana* needles or stems; M1, 2: synthesis of cDNA, 3: homogenizd cDNA, 4: post-column cDNA, 5: amplified plasmid, Lambda EcoT14 I digest (TaKaRa Dalian, China), M2. 250 bp DNA Ladder (TaKaRa Dalian, China).

**Fig S2. Construction of yeast two-bybrid Bait**

a: PCR amplication. According to WRKY31sequence, primers CLA354_WRKY31: CATGGAGGCCGAATTCATGGAAGCAGTGGGGTTGAGTCTT and CLA354_ WRKY31: GCAGGTCGACGGATCCTCACTGGAGCAATTTAGCCGAAGC were designed PCR amplication. b: Target fragment. c: Amplification of CLA354_WRKY31 plasmid as template. primersCLA420_WRKY31: CATGGAGGCCGAATTCATGGAAGCAGTGGGGTTGAGTCTT and, CLA420_WRKY31: GCAGGTCGACGGATCCTCACTGGAGCAATTTAGCCGAAGC were designed. d: Target fragment. M: 250bp DNA Marker.

**Fig S3. Bait strain self-activation test and two-hybrid screening**

a: Self-activation and toxicity assay of bait strain. Transformed bait plasmid into Y2Hgold strain to obtain bait strain, and bait strain was tested for self-activation and toxicity. b-d: Double-deficiency screening, Bait-Y2HGold strains were cultured for 3 days, and 2-3 mm colonies were picked up to incubate in SD/-Trp liquid medium, later, mating test was performed with a Y187 yeast library. Shake the bacteria slowly at 30-50 rpm for 24 h with incubation temperature 30 °C. Washed and suspended the yeast cells with 0.5 x YPDA (50 ug/mL kana). Diluted a small amount of the suspension by 1/10, 1/100, 1/1000, 1/10000 and applied 100 uL of the suspension to 100 mm monitor plates, and 50-55 SD/-Trp/-Leu/X-a-Gal/Aba double-deficiency screening plates were applied the suspension for screening.

**Fig S4. Vector construction of BiFC test for four genes**

Using needles and stems of *Pinus massoniana* as materials, amplification primers were designed according to the gene sequences of lp8, wrky2, wrky6 and wrky31. The target genes were amplified by using plasmids and cDNA as templates. The target gene lp8 was constructed on the pSPYNE-35S vector, and the wrky6, wrky2 and wrky31 were constructed on pSPYCE-35S vector.

**Fig S5. Defense reaction in of ABA and SA pathways with PmWRKY31**

a: Continuous variation of volatile matter content of needles in different treatments. b: Effects of different treatments on feeding and excretion of Dendrolimus. Each sample was repeated 3 times, * p< 0.05, ** p< 0.01, Student’s t-test.

**Table S1 Information on the primers in the experiment.**

**Table S2 Bioinformatics analysis of 4 genes and domain prediction.**

## Authors, Contributions

H. C. designed and conducted the experiments, and write the manuscript; X.L. contributed to manuscript writing and editing, Y.H. performed the bioinformatics tools, J.X., H.X., and Q.L. performed the physiology biochemistry experiment and analyzed of data, Z.Y. contributed to experimental design and editing. All authors read and approved the final manuscript.

